# Increased interhemispheric functional connectivity during non-dominant hand movement in right-handed subjects

**DOI:** 10.1101/2023.07.09.548228

**Authors:** Tomokazu Tsurugizawa, Ai Taki, Andrew Zalesky, Kazumi Kasahara

## Abstract

Hand preference is one of the behavioral expressions of lateralization in the brain. Previous fMRI studies, using conventional task-based functional MRI (fMRI), investigated the lateralization of brain function, and several regions including the motor cortex, putamen, thalamus, and cerebellum were activated during the single-hand movement. However, lateralization of functional connectivity related to hand preference has not been investigated. Here, we used the generalized psychophysiological interaction (gPPI) approach to investigate the alteration of functional connectivity during single-hand movement from the resting state in right-hand subjects. The functional connectivity in interhemispheric motor-related regions including the supplementary motor area, putamen, precentral gyrus, and cerebellum was significantly increased during non-dominant hand movement, while functional connectivity was not increased during dominant hand movement. The general linear model (GLM) showed activation in the left supplementary motor area, left precentral gyrus, and right cerebellum during right hand movement and activation in the right supplementary motor area, right precentral gyrus, and left cerebellum during left hand movement. These results indicate that a combination of GLM and gPPI analysis can detect the lateralization of hand preference more clearly in right-handed subjects.

## Introduction

Hand preference is one of the behavioral expressions of asymmetry in the human brain. Previous functional MRI (fMRI) studies, which rely on the blood oxygenation level-dependent (BOLD) signal change, investigated the lateralization of brain function. The task-based BOLD response is derived from changes in cerebral blood flow, cerebral blood volume (CBV), and the ratio of oxy/deoxy-haemoglobin, coupled with neuronal activation, which is evoked by psycho-physiological task or sensory input ^1^. A stimulus-induced signal change is calculated with the time course of the convolved hemodynamic response function with the timing of task ^2, 3^. Standard analysis of task-based fMRI data with general linear model (GLM) approach to separate stimulus-induced signal change from noise, which includes the fluctuation of the BOLD signals ^4^. Task-based fMRI with GLM during single-hand movement detects activation of regions, including the contralateral motor cortex, putamen, thalamus, and ipsilateral cerebellum ^5–7^. High-resolution fMRI reveals that information from the finger tapping is processed along closed-loop cortico-basal ganglia circuits ^8^. The use of the dominant hand was associated with a greater volume of activation in the contralateral motor cortex than that of the non-dominant hand, indicating the lateralization of the motor area ^9^. In addition to the regional activation in GLM analysis, the causality of BOLD response during simple hand movement was investigated with dynamic causal modeling (DCM). DCM only finds a difference in effective connectivity for different models if they predict sufficiently distinct patterns of BOLD responses ^10^. During unimanual movements, effective connectivity towards the contralateral primary motor cortex is enhanced, while effective connectivity towards ipsilateral motor areas is reduced by both transcallosal inhibition and top-down modulation ^11^. The contralateral sensorimotor area showed a stronger influence on the contralateral primary motor cortex when performing dominant hand movements ^12^. These results indicate that the activated area and stronger effective connectivity in the contralateral motor cortex indicates the lateralization of the brain associated with hand preference.

The functional connectivity, which reflects the functional network of the brain ^13^, is generally calculated from the fluctuation of the BOLD signals at the resting state that is task-free fMRI, from the correlation of the low-frequency fluctuation of the BOLD signals between anatomically separated regions ^14, 15^. The signal source of functional connectivity is not fully understood, but it is reported that spontaneous CBV fluctuation is correlated with spontaneous neuronal activity ^16^. The functional connectivity thus reflects the synchronization of the neuronal oscillation. The resting state fMRI can detect a wide range of large-scale brain networks associated with heterogenous cognitive processes ^17^ and it is increasingly being used to investigate the abnormal functional network in psychiatric disorders ^18^, neurodegenerative diseases ^19^, and developmental disorders ^20^. The functional connectivity, therefore, indicates distinct information such as neuronal synchronization compared with GLM analysis. To investigate the functional connectivity in task-based fMRI, another approach called the generalized psychophysiological interaction (gPPI), is required. The concept and framework of a PPI analysis are originally described by Friston and colleagues ^21^. BOLD response for task-based fMRI calculates the local activation, while gPPI shows the change in functional connectivity between anatomically separated regions affected during the task. In 2012, McLaren and colleagues applied this original framework for all task phases, and their interactions are included in a single model ^22^. This approach leads to results of interaction among the regions, and it is more sensitive and specific to the task phase. Together, GLM and DCM detect the local activation and synaptic efficacy based on the BOLD response, which is derived from local neuronal activation, during task. In contrast, gPPI can detect a modified brain functional connectivity, which reflects the synchronization of the neuronal oscillation, during the task. However, there has been no study that investigated the modified functional connectivity by single-hand movement. In the present study, using gPPI, we aimed to investigate the altered functional connectivity during dominant or non-dominant hand motion in right-handed subjects. The GLM analysis was also performed to assess that BOLD response during single hand movement corresponded to previous studies. The GLM analysis depicts the region and cluster size with a substantial increase in BOLD signals. The significant changes in the functional connectivity between the regions of interests (ROIs) was investigated using ROI-ROI gPPI analysis. Additionally, the voxel-based gPPI was then investigated with the seeds showing the significant change of functional connectivity by ROI-ROI gPPI analysis.

## Results

### Activation maps during the dominant or non-dominant hand movement by GLM analysis

The activation map during the left or right-hand movement, which was significant at P < 0.05 (FWE-corrected, N = 26), was first investigated with GLM analysis (Fig. 1 and 2). The lateralized activation was observed in many regions dependent on the single left or right hand. The dominant right hand movement evoked a BOLD response in the right cerebellum, the bilateral occipital lobe, the left preCG, the left thalamus, the left central operculum, the left supplementary motor area (SMA), and the left putamen (Table 1 and Fig. 2A). During the non-dominant left-hand movement, a significant BOLD response was observed in the bilateral cerebellum, the bilateral occipital lobe, the right preCG, the right thalamus, the right central operculum, and the right SMA (Table 2 and Fig. 2B). The cluster size of the activated area that was significantly increased during the task from the group-level analysis was compared. The cluster size of the activated area in the contralateral preCG, and ipsilateral cerebellum during right hand movement was greater than that of non-dominant left-hand movement (Tables 1 and 2). The significant cluster size was calculated in each subject following first level analysis (p < 0.05, FWE-corrected) and statistically compared between right and left-hand movement (Table 3). There was no significant difference in the contralateral preCG and in the ipsilateral cerebellum (p > 0.05, paired t-test). The BOLD response in the bilateral occipital lobe, which relates to visual processing, was observed during left and right-hand movement. The statistical comparison of BOLD response (p < 0.05, FWE-corrected, N = 26) between left and right-hand movement showed that BOLD response in the contralateral preCG, and ipsilateral cerebellum was significantly increased in single-hand movement (Fig. 2C, Tables 4 and 5). Furthermore, the ipsilateral central operculum (CO) was significantly activated in right-hand movement only. These results indicate the activated regions of the brain regions regulating the motor function were almost symmetrical, such as the contralateral preCG, SMA, thalamus, putamen, and ipsilateral cerebellum, evoked by the single-hand movement. However, the larger cluster size of the activated area and a significant increase in the contralateral CO during dominant hand movement indicates the lateralization of single hand movement in the right-handed.

**Figure 1.**
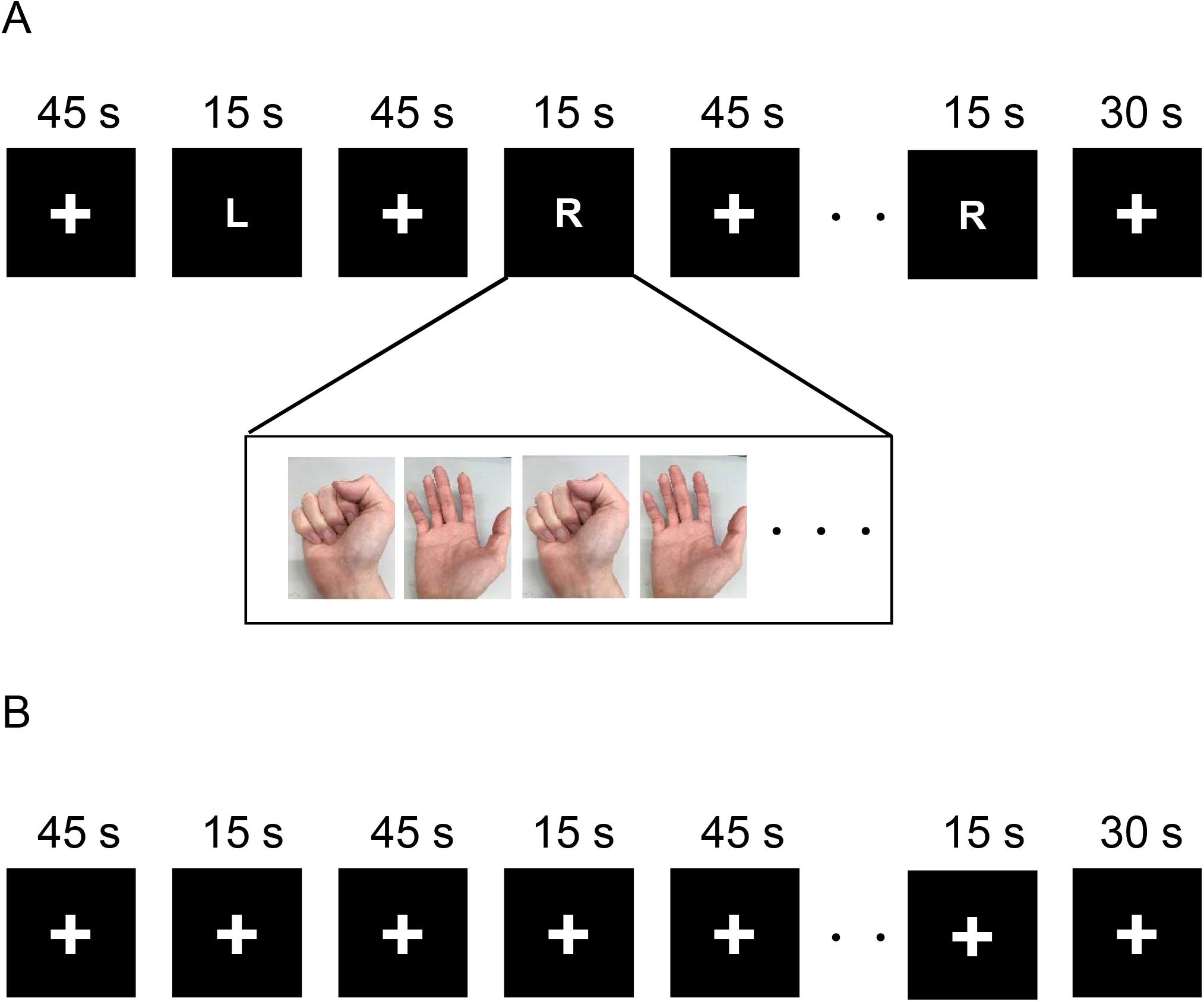
Schematic figure of the experimental paradigm. (A) The schematic figure of single hand movement experiment. The “+” indicates the rest period. The “L” means the left hand movement and the “R” means the right hand movement. During the single hand movement task, the subjects continued to open and close their fingers. The fMRI scanning was continued for 10 minutes and 30 seconds. (B) The schematic figure of fMRI without single hand movement.

**Figure 2.**
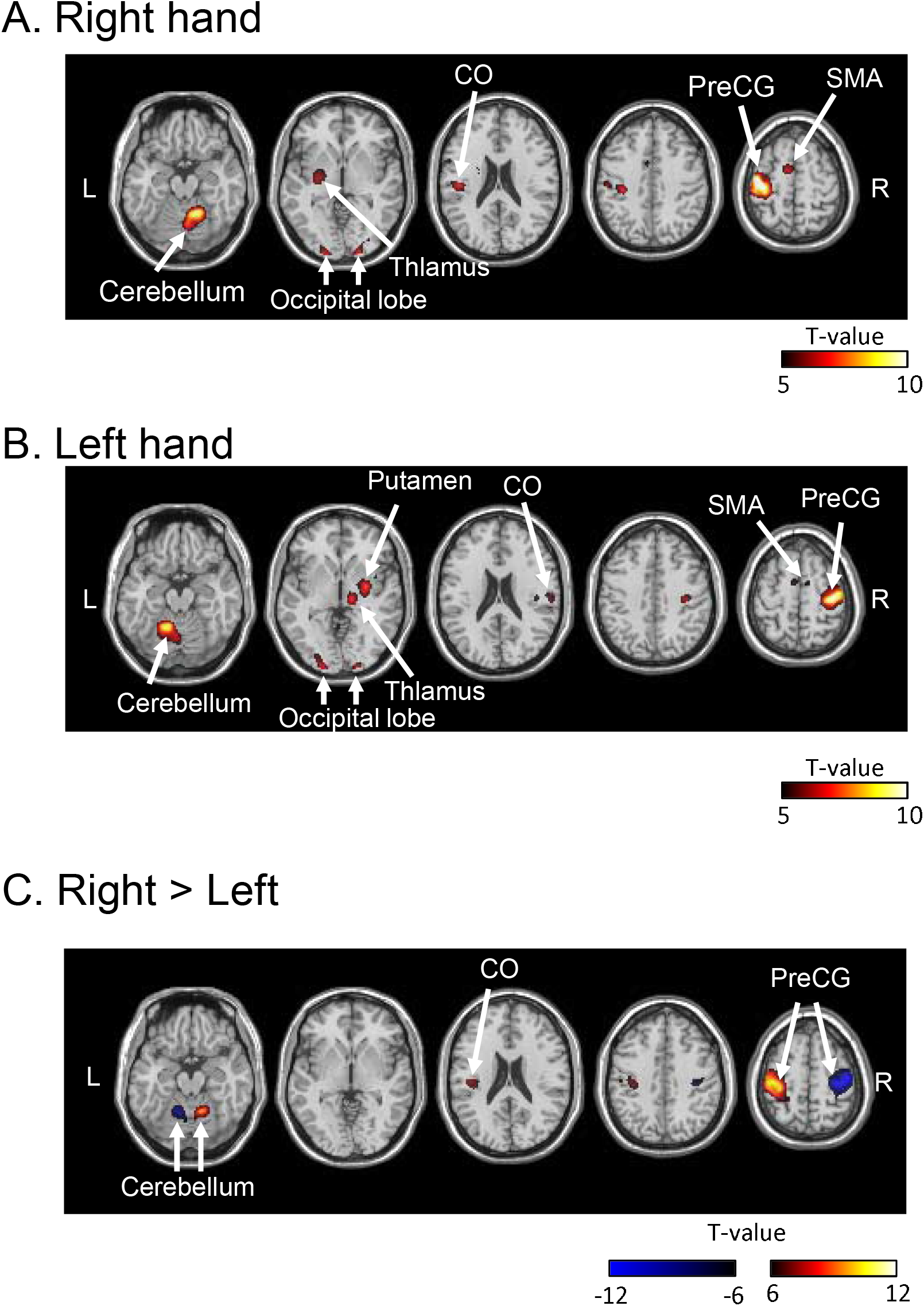
Significant increase of BOLD signal during single hand movement. (A) Significant BOLD increase during the right hand movement (p <0.05, family wise error (FWE)-corrected, N = 26). (B) The significant BOLD increase during the left hand movement (p < 0.05, FWE-corrected, N = 26). (C) Significant difference between right hand movement and left hand movement (p<0.05, FWE-corrected, N = 26). L, left; R, right. The color bar indicates the t-values. CO, central operculum; preCG, precentral gyrus; SMA, supplementary motor area.

**Table 1.**
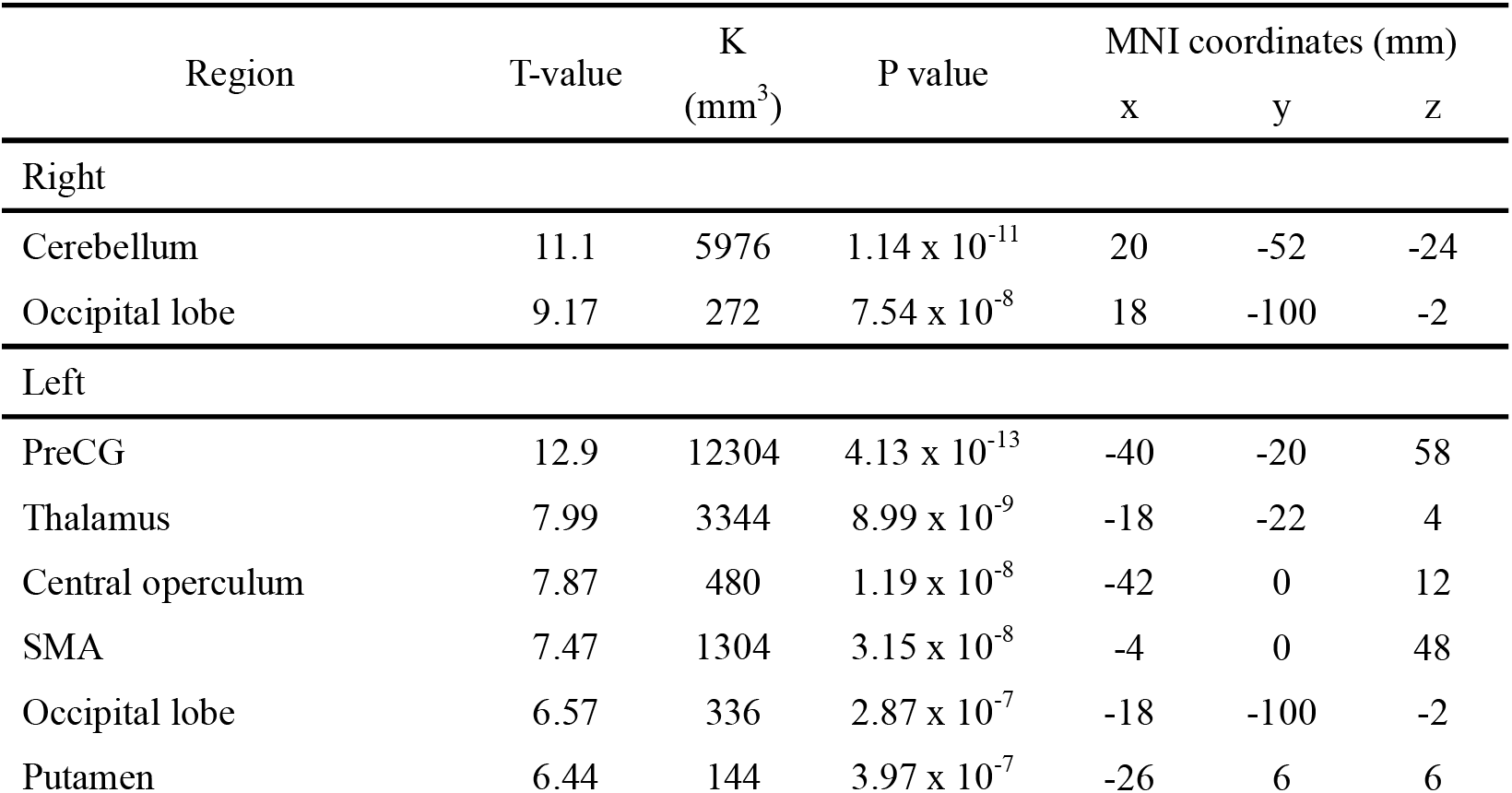
The significant BOLD increase during the dominant right-hand movement. T-value indicates the peak-level t-value. K indicates the cluster size. P-value indicates peak-level p-value. Right and left indicate each hemisphere.

**Table 2.** The significant BOLD increase during the non-dominant left-hand movement. T-value indicates the peak-level t-value. K indicates the cluster size. P-value indicates peak-level p-value. Right and left indicate each hemisphere.

| Region | T-value | K<br>(mm<sup>3</sup>) | P value | MNI coordinates (mm) |  |  |
| --- | --- | --- | --- | --- | --- | --- |
|  |  |  |  | x | y | z |
| Right |  |  |  |  |  |  |
| PreCG | 13.5 | 10912 | 1.50 x 10<sup>-13</sup> | 38 | -18 | 52 |
| Thalamus | 9.17 | 3384 | 6.22 x 10<sup>-10</sup> | 12 | -20 | 2 |
| Central operculum | 7.59 | 492 | 2.34 x 10<sup>-8</sup> | 46 | -16 | 16 |
| Occipital lobe | 7.46 | 152 | 3.22 x 10<sup>-8</sup> | 20 | -98 | -2 |
| Cerebellum | 7.07 | 88 | 8.31 x 10<sup>-8</sup> | 26 | -60 | -22 |
| SMA | 15 | 120 | 5.22 x 10<sup>-7</sup> | 62 | -22 | 26 |
| Left |  |  |  |  |  |  |
| Occipital lobe | 8.45 | 96 | 1.89 x 10<sup>-8</sup> | -22 | -96 | 2 |
| Cerebellum | 12.1 | 4632 | 1.89 x 10<sup>-8</sup> | -16 | -52 | -20 |

**Table 3.** Averaged cluster size of the significant increase in each subject during left and right hand movement. Data are expressed as mean ± SEM (mm^3^).

|  | PreCG |  | Cerebellum |  |
| --- | --- | --- | --- | --- |
|  | Left | Right | Left | Right |
| Right-hand | 34677 ± 7034 | - | - | 14290 ± 3416 |
| Left-hand | - | 23781 ± 4039 | 10052 ± 2332 | - |

**Table 4.**
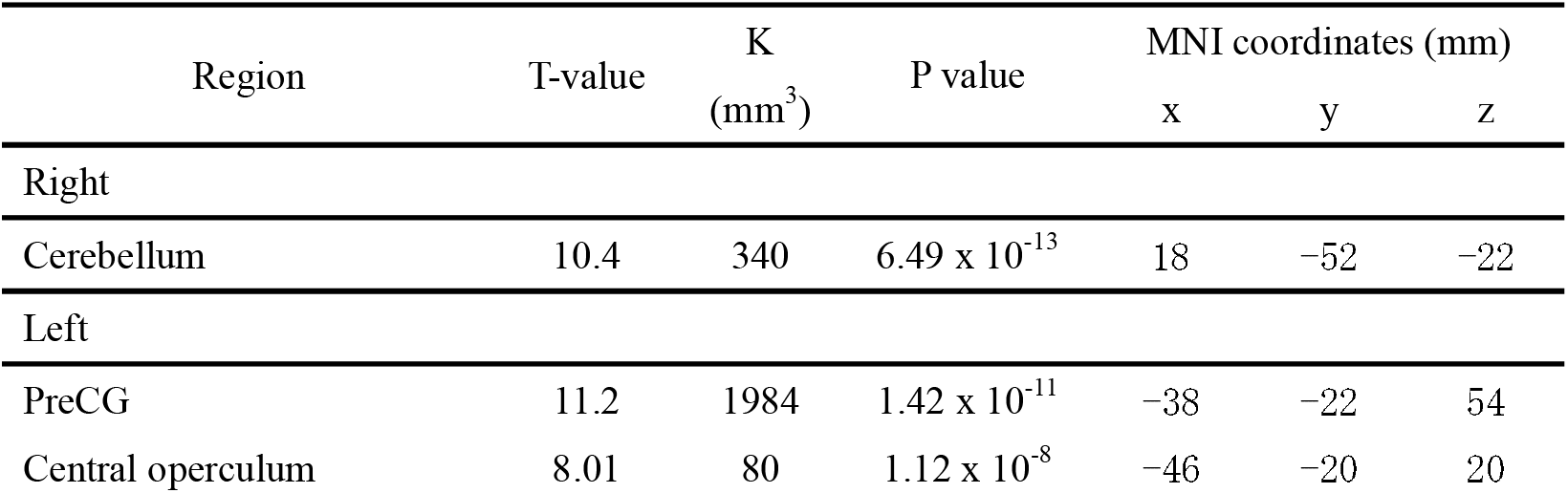
The significant BOLD increase during the dominant right-hand movement compared with left hand movement. T-value indicates the peak-level t-value. K indicates the cluster size. P-value indicates peak-level p-value. Right and left indicate each hemisphere.

The bilateral BOLD response in the visual cortex was evoked during single hand movement, corresponding to the previous study, which reported the activation of the occipital lobe by single-letter stimulation ^23^. These activations were not different between right hand and left-hand movement (Fig. 2C). Furthermore, as the negative control, the BOLD signal change without single hand movement task was assessed. The indicator of the left- and right-hand movement was not presented, and participants did not move their hands (Fig. 1B). The same regressor as the single-hand movement was used for GLM analysis, and the BOLD response was not observed. These results support that the observed BOLD response was caused by the single hand movement.

### ROI-ROI gPPI

Figures 3 and 4 show the significant increase or decrease of functional connectivity during single hand movement (p < 0.05, NBS-corrected, N = 26). A significant increase of interhemispheric functional connectivity was observed in many regions during the non-dominant left hand movement (Fig. 3A). From these strengthened connections, the significant changes in functional connectivity between the regions associated with motor regulation are focused on Fig. 3B. In the right hemisphere, the putamen, the SMA, the inferior frontal gyrus operculum (IFGoper), the anterior/posterior supramarginal gyrus (aSMG and pSMG), the preCG and the postcentral gyrus (postCG) increased the functional connectivity with many bilateral regions. In the left hemisphere, the putamen and the SMA significantly increased the functional connectivity with the multiple regions in the right hemisphere. Especially, the right preCG, the left SMA, the right IFGoper, the right aSMG, the right pSMG, and the right putamen significantly increased the functional connectivity with more than five brain regions (Table 5). A significant decrease in functional connectivity was also observed in the frontal medial cortex, bilateral cerebellum/vermis, and the left anterior middle temporal gyrus (aMTG) (Fig. 3C). During the right-hand movement, a significant increase in functional connectivity was not observed, while a few functional connections including the bilateral caudate, the left thalamus, and the right cerebellum were decreased (Fig. 4A). The direct comparison of functional connectivity between left and right-hand movement was then performed (Fig. 4B). The increased functional connectivity during left-hand movement rather than right-hand movement was observed in the bilateral brain regions including the bilateral SMA, the right IFGoper, the left postCG, and the bilateral thalamic nuclei.

**Figure 3.**
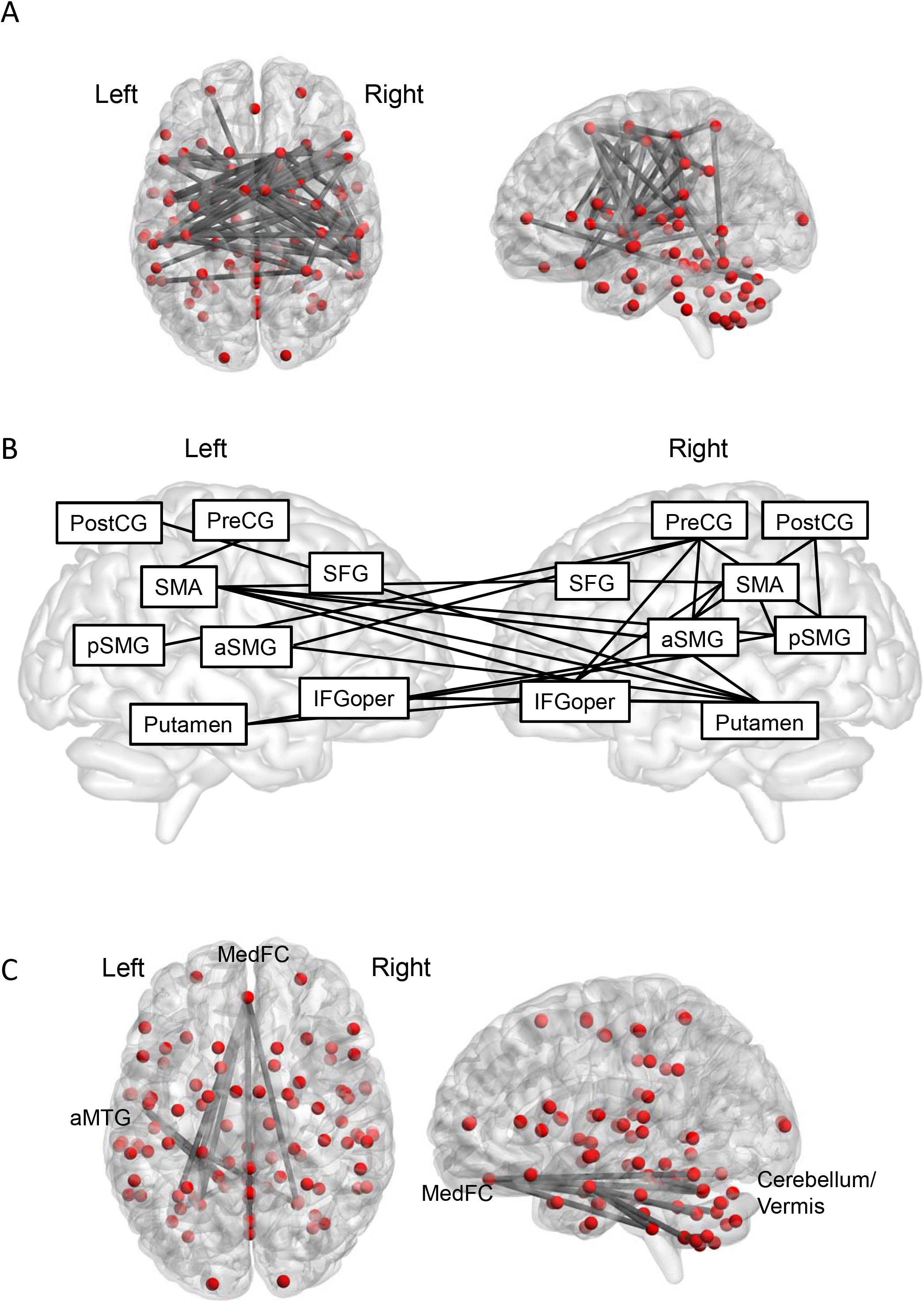
Altered functional connectivity during the single left-hand movement. (A) Brain and circle plots for all nodes during the left hand movement. The Grey line indicates the increased connectivity (p<0.05, NBS-corrected, N = 26). (B) Significantly increased functional connectivity in the motor network during left hand movement (p<0.05, NBS-corrected, N = 26). The Red line indicates the connectivity with the right inferior frontal gyrus operculum (IFGoper). The green line indicates the connectivity with the right precentral gyrus (preCG). The yellow line indicates the connectivity with the right supplementary motor area (SMA). The blue line indicates the connectivity with the right putamen. The purple line indicates the connectivity with the left SMA. (C) Brain and circle plots for all nodes during the left hand movement. The Grey line indicates the significantly decreased connectivity (p < 0.05, NBS-corrected, N = 26).

**Figure 4.**
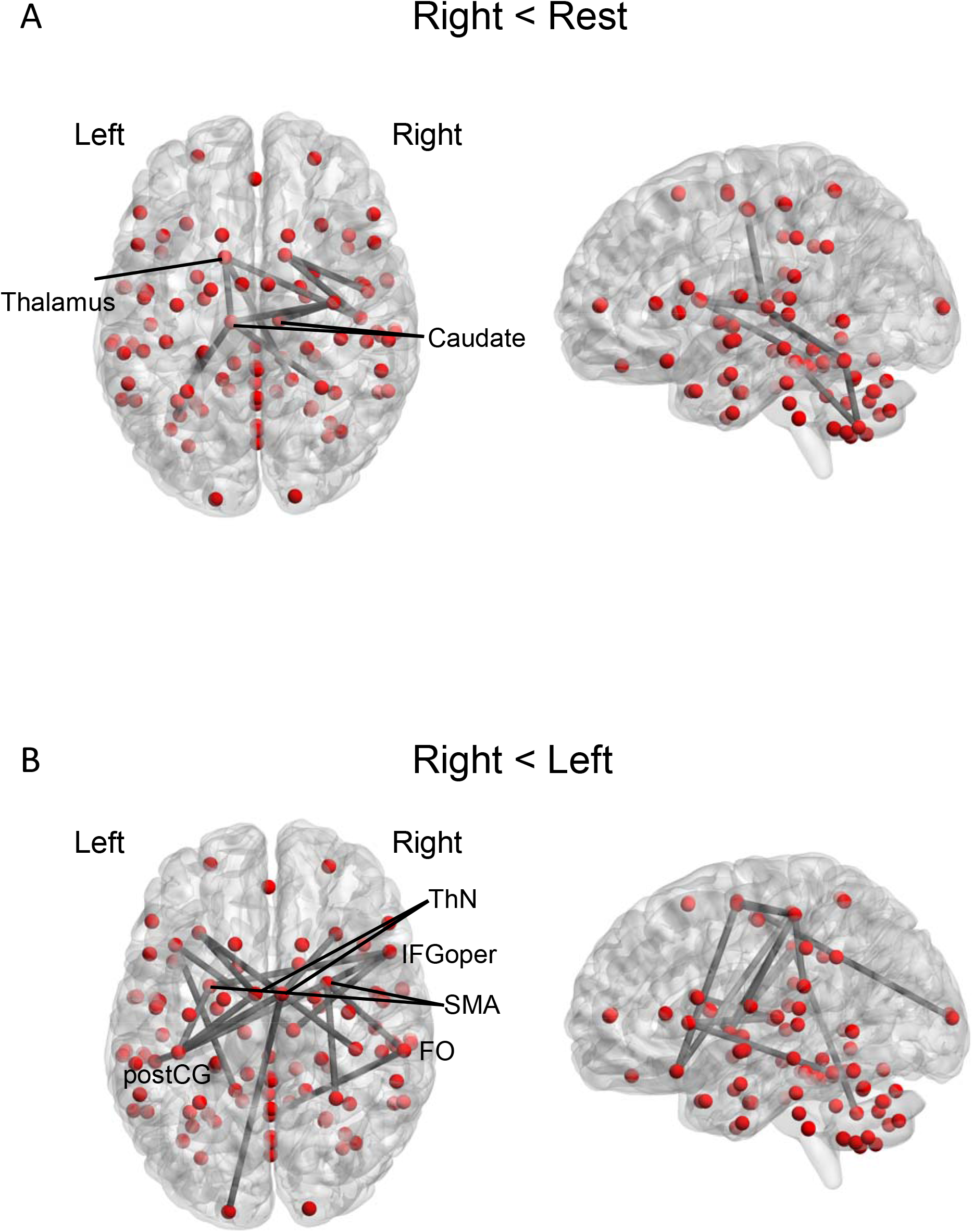
Altered functional connectivity during the single right-hand movement. (A) Brain and circle plots for all nodes during the right-hand movement. The Grey line indicates the significantly decreased functional connectivity (p<0.05, NBS-corrected, N = 26). (B) Significant increase of functional connectivity during left-hand movement rather than right-hand movement (p < 0.05, NBS-corrected, N = 26).

**Table 5.** The significant BOLD increase during the non-dominant left-hand movement compared with right hand movement. T-value indicates the peak-level t-value. K indicates the cluster size. P-value indicates peak-level p-value. Right and left indicate each hemisphere.

| Region | T-value | K<br>(mm<sup>3</sup>) | P value | MNI coordinates (mm) |  |  |
| --- | --- | --- | --- | --- | --- | --- |
|  |  |  |  | x | y | z |
| Right |  |  |  |  |  |  |
| PreCG | 12.1 | 1799 | 3.13 x 10<sup>-12</sup> | 30 | -22 | 60 |
| Left |  |  |  |  |  |  |
| Cerebellum | 9.46 | 394 | 4.82 x 10<sup>-10</sup> | -18 | -52 | -22 |

The change of functional connectivity without single hand movement was also assessed (Fig. 1B). The method of gPPI analysis was same as that with single hand movement, that is, same psychological term for single hand movement task was used, and a significant increase or decrease in functional connectivity was not observed. These results support that the observed changes in functional connectivity were caused by the single hand movement.

### Seed-based gPPI

The significant change in functional connectivity during single hand movement was also investigated at the voxel level (p < 0.05, NBS-corrected, N = 26). The seed was decided from Table 6, such as the left SMA, the right IFGoper, the right preCG, the right putamen, the right aSMG, and the right pSMG. The functional connectivity with the left SMA increased in the bilateral insular cortex, the right preCG, the right superior temporal gyrus (STG), the right temporooccipital part of MTG (toMTG), and the left putamen (Fig. 5A). The functional connectivity with the right IFGoper increased in the anterior part of the bilateral putamen, the bilateral IC, the bilateral SMA, the bilateral superior frontal gyrus (SFG), and the right preCG (Fig. 5B). The functional connectivity with the right preCG increased in the anterior part of the bilateral medial frontal gyrus (MFG), the bilateral anterior sensorimotor gyrus (aSMG), the right IFGoper, and the left frontal operculum (Fig. 5C). The functional connectivity with the right putamen increased in the bilateral IC, the bilateral IFGoper, the bilateral precentral gyrus medial segment (MPrG), the left planum temporale (PT), and the left preCG (Fig. 5D). The functional connectivity with the right aSMG increased in the bilateral MPrG, the bilateral SMA, and the right preCG (Fig. 5E). The functional connectivity with the right pSMG increased in the bilateral SMA, the bilateral MrPG, the right preCG, the left Heschl’s gyrus (HG), and the left PT (Fig. 5F). We did not observe any change of the functional connectivity with these regions during the right-hand movement. These six ROIs showed the significant increase of interhemispheric functional connectivity during non-dominant left-hand movement, not during dominant right hand movement, corresponding to those by ROI-ROI analysis.

**Figure 5.**
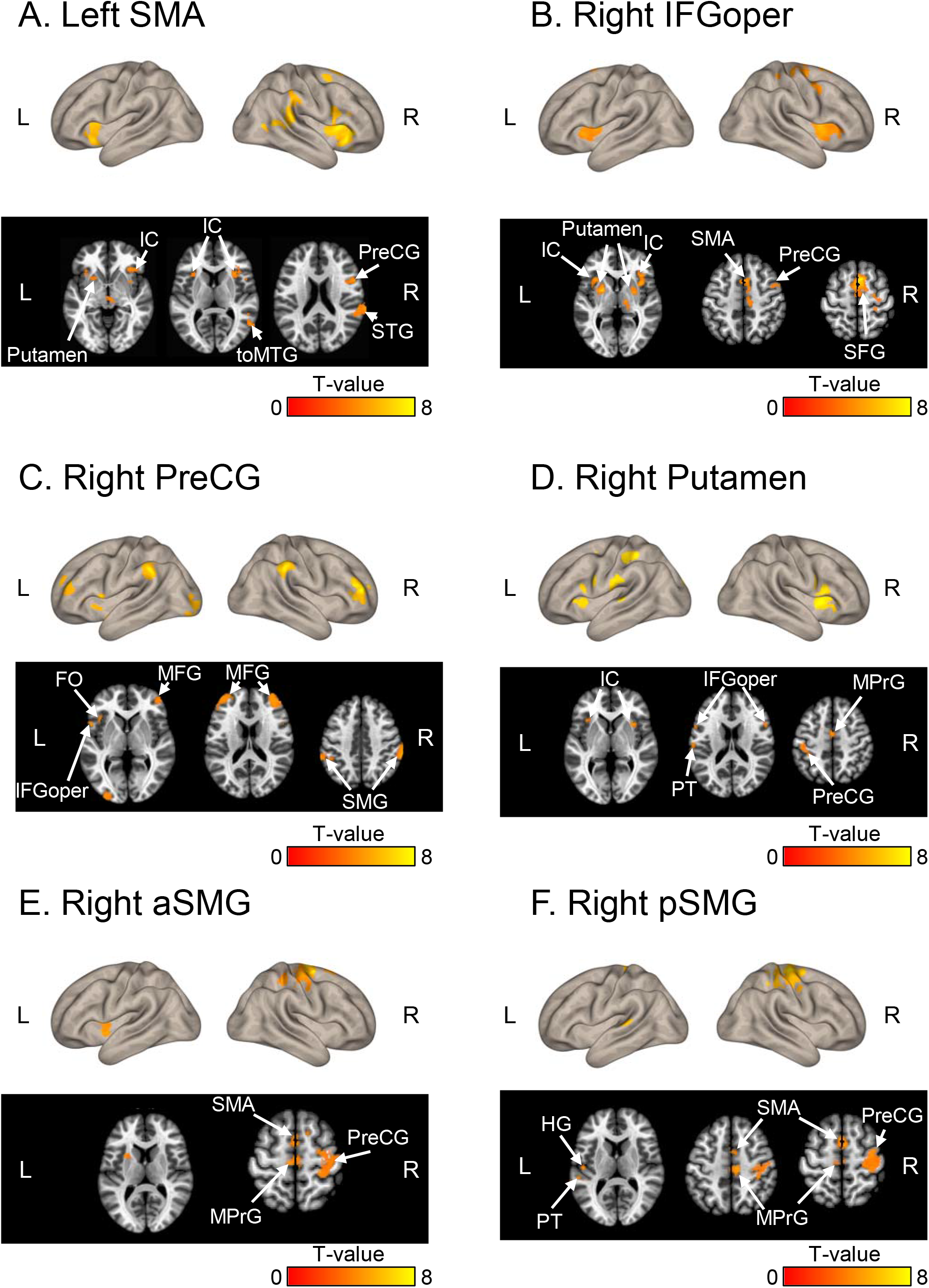
Voxel-based gPPI analysis. Increased functional connectivity (A) with the left supplementary motor area (SMA), (B) with the right inferior frontal gyrus operculum (IFGoper), (C) with the right precentral gyrus (preCG), (D) with the putamen, (E) with the right anterior supramarginal gyrus (aSMG), and (F) with the posterior supramarginal gyrus (pSMG) during left hand movement. P< 0.05, FEW-corrected, N = 26. Color bar, T-values. FO, frontal operculum; FP, frontal pole; IC, insular cortex; IFGoper, inferior frontal gyrus operculum; MFG, middle frontal gyrus; MPrG, Precentral gyrus medial segment; HG, Heschl’s gyrus; OP, occipital pole; PT, planum temporale; SFG, superior frontal gyrus; SMG, superior middle gyrus; STG, superior temporal gyrus; toMTG, temporooccipital part of middle temporal gyrus.

**Table 6.** The regions having more than five connections those significantly increased functional connectivity during non-dominant left hand movement. P-value indicates peak-level p-value.

| Seed Region | Significantly increased connectivity<br>($p < 0.05$, NBS-corrected) |
| --- | --- |
| Left SMA | left preCG, right SFG, right IFGoper, right putamen, right aSMG, right pSMG |
| Right IFGoper | Left/right SMA, left/right putamen, right preCG |
| Right preCG | Left SMA, left/right pSMG, right IFGoper, right aSMG |
| Right putamen | Left postCG, left SMA, left/right aSMG, right IFGoper |
| Right aSMG | Left/right SMA, left/right putamen, right preCG, right postCG |
| Right pSMG | Left/right SMA, left putamen, right preCG, right postCG |

The change of functional connectivity without single hand movement was also assessed (Fig. 1B). The method of gPPI analysis was same as that with single hand movement, that is, same psychological term for single hand movement task was used, and a significant increase or decrease in functional connectivity was not observed. These results support that the observed changes in functional connectivity were caused by the single hand movement.

## Discussion

In the present study, we successfully showed a change in the functional connectivity during single dominant/non-dominant hand movement compared with that in the rest period. The classical GLM analysis showed activation in the contralateral supplementary motor area, contralateral precentral gyrus, and ipsilateral cerebellum during hand movement. (Fig. 2). In contrast, the activated area in the contralateral SMA and preCG during dominant right hand movement was greater rather than that during non-dominant left hand movement, consistent with the previous study ^9^. In contrast to GLM analysis, gPPI showed interhemispheric functional connectivity, including the primary motor regions, such as the putamen, the preCG, and the SMA during non-dominant hand movement (Figs. 3-5).

The gPPI enables us to investigate the altered functional connectivity during the psycho-physiological task ^24–26^. In the present study, using gPPI, we clearly showed asymmetric connectivity between the dominant and non-dominant hand movement. The non-dominant hand movement significantly increased the interhemispheric functional connectivity in many brain regions centered in the motor network from the resting state rather than the dominant hand movement. The different results between GLM and gPPI analysis are possibly explained by the distinct information of BOLD signals. The GLM focuses on the BOLD signal response evoked by the task, that is an increase of BOLD signal intensity triggered by the neuronal activation, while gPPI focuses on the change of the correlation of fluctuation of psychophysiological term evoked by the task. The previous study also demonstrated that local activation was not related to connectivity changes in large scale brain networks, corresponding to results of current study ^27^. The local brain activation and task dependent connectivity change can be modulated in different directions at the same time. This can be explained by the neuronal mechanism of fMRI signal changes. The BOLD response is coupled with the local neuronal activity ^28^, while functional connectivity is related to the coordinated interaction of excitatory and inhibitory inputs as well as the irregular firing of single neurons ^29^. Although the frequency range between electroencephalography (EEG) (0.1 – 100 Hz) and fMRI (0.008 – 0.08 Hz) is different, the previous EEG-fMRI studies show that functional connectivity is correlated with the synchronization of not only gamma oscillation ^30, 31^ but also alpha and beta oscillation of the local field potential ^32^. By simultaneous EEG-fMRI, in participants at rest, it has been shown that fMRI activity in brain networks was correlated with power fluctuations of neuronal oscillations, primarily in the alpha, beta, and gamma bands ^33^. The phase synchrony in the alpha and beta bands contributes particularly strongly to the functional connectivity calculated by fMRI. The altered functional connectivity by gPPI, therefore, results from the altered neuronal interactions associated with hand movement ^34^.

The BOLD signal increase in the posterior putamen and the sensorimotor cortex, preCG, and the cerebellum during the hand movement has been reported in the previous study ^8^, consistent with our results (Fig. 2). The use of the dominant hand was associated with a greater volume of activation in the contralateral motor cortex ^9^. Furthermore, the causality of the regional activation in motor regulation was investigated by the DCM approach. DCM is a method that has been successfully used to infer directed connectivity between brain regions based on task-based fMRI and possibly estimates of intrinsic synaptic coupling strengths ^35^. The dominant right-hand movement in right-handler shows significant coupling of contralateral SMA with ipsilateral SMA, contralateral putamen and contralateral motor area ^12^. Moulton et al investigated the effective connectivity among eight regions of interest in the visuomotor network (corticocortical and cerebello-cortical) for the control of grip force of the dominant vs. non-dominant hand in right-handers ^36^. They showed a coupled cerebellar-SMA and cerebellar-posterior parietal cortex modulation (bidirectional, i.e., feed-forward and feedback) specifically during visuomotor force control with the non-dominant hand movement. The altered effective connectivity was not observed during the dominant hand movement. Together, the effective connectivity by DCM shows the asymmetrical effective connectivity between the dominant and non-dominant hand movement differently with GLM analysis. In contrast to the DCM, this study shows a different kind of asymmetry of functional connectivity in hand movement. Remarkably, no significant change in functional connectivity was observed in dominant hand movement.

In the current study, the interhemispheric brain network centered on the motor network, including the contralateral SMA, preCG, the contralateral IFGoper, the contralateral putamen, and the ipsilateral cerebellum, were significantly increased during non-dominant hand movement (Fig. 3). Importantly, the left contralateral SMA increased the interhemispheric functional connectivity of the motor loop, including the ipsilateral putamen, SMG, and IFGoper, as well as with the contralateral preCG (Table 6). Previous EEG studies showed interhemispheric connectivity during non-dominant hand movement. The left-hand use facilitates the frontal N30, related to the activity in the basal ganglia, SMA, and primary motor area, somatosensory evoked potential ^37^. This indicates that coordination and synchronization between involved brain regions during non-dominant hand movement can be possibly caused by altered activity related to sensory gating. The interhemispheric functional connectivity in the Parietal-Frontal cortices, including the connectivity between the right superior parietal lobule and left/right primary motor cortices, is essential for non-dominant left-hand movement ^38^. In addition to the contribution of the contralateral motor cortex and basal ganglion to motor regulation of the limbs, there is increasing evidence that the ipsilateral hemisphere on the limb is also active during movements ^39^. Together with present studies, increased interhemispheric connectivity is related to the movement of the non-dominant hand. The previous studies using EEG investigated the part of functional connectivity during non-dominant hand movement and the detailed change of functional connectivity in the whole brain has not been shown. This is because EEG has low spatial resolution. The present study, using gPPI analysis, clearly shows the increased interhemispheric functional connectivity in the whole brain.

### Perspectives and Conclusions

The interhemispheric functional connectivity is increased during the non-dominant hand movement. In contrast, interhemispheric functional connectivity is not changed during dominant hand movement. Because GLM and gPPI show distinct phenomena of neuronal function, such as the local activation and synchronization of neuronal oscillation between the brain regions, studies in combination with GLM and gPPI analyses will provide new insight into the functional lateralization of our brain in humans.

### Limitation of study

Traditionally, left-handed people have been corrected to be right-handed and the percentage of left-handed is around 3% in Japanese ^40^. Therefore, we could not find enough subjects to compare with right-handed. In the future, we will plan to investigate left-handed to validate the laterality of single-hand movement, although it will take longer time to collect the left-handed participants than right-handed participants. In this study, the volunteers were not trained for the movement, so repeated training can change the functional connectivity during non-dominant hand movement.

In this study, we investigated fMRI in 26 subjects. Although previous studies showed the good quality of the results and the results of BOLD increased area are replicable with less than 30 subjects ^7, 36, 41^, a larger sample size would be preferable and is expected for simple fMRI studies ^42^.

The start of left-or right-hand movement was not randomized. Therefore, there might be the possibility that sequence bias exists, although this bias seems small because hand movement in each hand was performed five times alternatively.

### STAR Method Participants

Twenty-six healthy young adults (17 males and 9 females, ages 30 ± 1) participated in the fMRI experiment. All individuals were right-handed. Individuals were healthy, with no history of alcohol abuse, neurological disease, or learning disability. Right-handedness was determined using the laterality quotient from the 10-item version of the Edinburgh Handedness Inventory (EHI) ^43^ (Fig. S1), because some items in 20 item version, such as cricket bat, are not familiar with Japanese. The laterality quotient is calculated as (R-L)/(R+L) × 100, where R and L is the number marked on the right and the left side respectively. The right-handed participants were eligible for inclusion with EHI scores > 40 by EHI, which is based on previous studies ^43, 44^. All experimental procedures and protocols were approved by the Institutional Review Board of the National Institute of Advanced Industrial Science and Technology.

### fMRI task and acquisition parameters

MRI images were acquired using a 3T Ingenia MRI system with a 32ch brain coil (Philips, Netherlands). The structural image was acquired by magnetization prepared rapid gradient echo (MPRAGE) as follows: TE / TR = 5.1 / 11 ms, flip angle = 8 degrees, matrix = 368 × 315 × 44, resolution = 0.70 × 0.76 × 0.70 mm^3^ / voxel. The fMRI was acquired to assess functional connectivity during hand motion with the following parameters: TE / TR = 30 / 1,500 ms, flip angle = 80 degrees, matrix = 76 × 76 × 44, resolution = 2.5 × 2.5 × 2.5 mm^3^ / voxel. The tasks were presented as a block design, which contains the 5 left hand movements and 5 right hand movement blocks (15 s) alternatively interpreted with a rest period (45 s) (Fig. 5). The single hand task started from the left hand. During the hand movement period, subjects repeated to open and close hands by moving all fingers and the hands were moved, neither up nor down. The rest period, at last, was the 30s; scanning continued for 10.5 min. During the rest period, participants watched the cross mark on the screen. The participants were indicated to close and open their hands around 1-1.5 Hz because the previous study reveals that the amplitude of the BOLD response is a plateau for hand movement rates larger than 1 Hz^41^.

As a negative control, the subjects performed fMRI without single hand movement (Fig. 1B). The “+” mark was presented during the single hand movement task. The fMRI scan was continued for 10.5 min. The observed data were analyzed statistically with the same protocol as single hand movement.

### Preprocessing of fMRI data

Statistical parametric mapping SPM12 software (Welcome Trust Center for Neuroimaging, UK) was used to analyze the fMRI data and to perform preprocessing steps, including slice timing correction, motion correction by realignment, normalization, and smoothing with a Gaussian filter (8 × 8 × 8 mm²/voxel of half-width at half-maximum). For connectivity analysis, pre-processed fMRI data were then detrended, and slow periodic fluctuations were extracted using a bandpass filter (0.008–0.08 Hz). A total of 96 regions of interest (ROIs), which are from the automated anatomical labeling atlas, were used for ROI analysis. The correlation coefficients between two ROIs were calculated using the CONN toolbox.

### BOLD response analysis

For the first level (fixed effect analysis), an onset regressor defined the onsets of left or right blocks, and the block length was set to 15 s for each session. The hemodynamic response was modeled using a canonical hemodynamic response function in SPM12. The six head movement parameters, which were estimated in the realignment process, were included as regressors to regress out the effect of head motion. The resulting contrast images were calculated at first level by setting the targeting regressor (regressor of left or right-hand motion) to 1 and others to zero. The significant cluster size (p < 0.05, FEW-corrected) in the preCG and cerebellum was calculated in each subject following first level analysis. These contrasts were then used for a second-level analysis (group analysis). For the second level, model estimation was performed using the one-sample t-test as implemented in SPM12. The significant voxels were thresholded at P < 0.05, FWE-corrected.

### ROI-to-ROI gPPI

The ROI-to-ROI generalized psychophysiological interaction (gPPI) was performed with CONN toolbox (www.conn-toolbox.org) to investigate which brain regions interact in a task-dependent manner ^22^. The gPPI analysis detects the brain regions whose connectivity with another ROI varies as a function of psychological context, that is, left or right-hand motion in this experiment. The 96 ROIs in the right and left hemispheres were defined in the previous sections. ROI-to-ROI gPPI computation was made for every possible pair-wise combination of selected ROIs for each individual. Predictors include the estimated time course of the task (psychological term), the time-series regressor of BOLD signal changes in each ROI (physiological term), and interactions between psychological and physiological terms (PPI term), which consists of the product of the psychological term multiplied by the physiological term. For the first-level analysis, a PPI regressor (PPI term) was generated for each condition (rest, left hand, or right hand) as a product of the ROI time series multiplied by the task effect, and the beta weight was calculated in all ROIs. At the group level, random effect analysis was used across participants, and the one-sample t-test was calculated to compare ROI-based connectivity for rest vs. left, rest vs. right, or left vs. right conditions.

The statistical significance of ROI-ROI connectivity was assessed by paired t-test with a corrected threshold of p < 0.05, using the Network-Based Statistic toolbox (NBS; https://www.nitrc.org/projects/nbs/). The primary threshold for the NBS was set to a Cohen’s d of 0.7 (minimum meaningful effect size). Subnetworks were then identified in the set of supra-threshold connections. The size of each subnetwork was measured in terms of the number of connections it comprised. Permutation testing was used to compute a family-wise error-corrected p-value for each subnetwork. Specifically, 5000 permutations were generated in which group labels were randomly permuted and the size of the largest subnetwork was recorded for each permutation. Subnetworks deemed to be significant with permutation testing (p < 0.05, FWE-corrected) were identified as significant.

### Seed-to-voxel gPPI

For seed-to-voxel gPPI, an averaged BOLD time course within selected ROI was extracted for each seed-ROI and used as a physiological regressor in each individual using CONN toolbox. For the first-level analysis, a PPI regressor (interaction term) was generated for each condition as the product of the ROI time series multiplied by the psychological term (boxcar function of rest, left-hand, or right-hand cues). The seed was the regions having more than five connections those significantly increased functional connectivity during non-dominant left hand movement from ROI-ROI gPPI analysis, that is, the left SMA, the right IFGoper, the right preCG, the right putamen, the right aSMG, and the right pSMG. Significant voxels were determined by p < 0.05, FWE-corrected.

## Acknowledgments

We thank Ms. Natsu Nozaki for the support of the MRI experiment.

## Data and code availability statement

The data supporting the reported findings are available from the corresponding author upon reasonable request.

## Author contribution

**Tomokazu Tsurugizawa:** Conceptualization, Methodology, Investigation, Data Analysis, Visualization, Writing - review & editing. **Ai Taki:** Investigation, Data Analysis, Writing - review & editing. **Andrew Zalesky:** Data analysis, Visualization, Writing - review & editing. **Kazumi Kasahara:** Conceptualization, Methodology, Writing - review & editing.

